# Transposable elements contribute to activation of maize genes in response to abiotic stress

**DOI:** 10.1101/008052

**Authors:** Irina Makarevitch, Amanda J. Waters, Patrick T. West, Michelle C. Stitzer, Jeffrey Ross-Ibarra, Nathan M. Springer

## Abstract

Transposable elements (TEs) account for a large portion of the genome in many eukaryotic species. Despite their reputation as “junk” DNA or genomic parasites deleterious for the host, TEs have complex interactions with host genes and the potential to contribute to regulatory variation in gene expression. It has been hypothesized that TEs and genes they insert near may be transcriptionally activated in response to stress conditions. The maize genome, with many different types of TEs interspersed with genes, provides an ideal system to study the genome-wide influence of TEs on gene regulation. To analyze the magnitude of the TE effect on gene expression response to environmental changes, we profiled gene and TE transcript levels in maize seedlings exposed to a number of abiotic stresses. Many genes exhibit up- or down-regulation in response to these stress conditions. The analysis of TE families inserted within upstream regions of up-regulated genes revealed that between four and nine different TE families are associated with up-regulated gene expression in each of these stress conditions, affecting up to 20% of the genes up-regulated in response to abiotic stress and as many as 33% of genes that are only expressed in response to stress. Expression of many of these same TE families also responds to the same stress conditions. The analysis of the stress-induced transcripts and proximity of the transposon to the gene suggests that these TEs may provide local enhancer activities that stimulate stress-responsive gene expression. Our data on allelic variation for insertions of several of these TEs show strong correlation between the presence of TE insertions and stress-responsive up-regulation of gene expression. Our findings suggest that TEs provide an important source of allelic regulatory variation in gene response to abiotic stress in maize.

## Author summary

Transposable elements are mobile DNA elements that are a prevalent component of many eukaryotic genomes. While transposable elements can often have deleterious effects through insertions into protein-coding genes they may also contribute to regulatory variation of gene expression. There are a handful of examples in which specific transposon insertions contribute to regulatory variation of nearby genes, particularly in response to environmental stress. We sought to understand the genome-wide influence of transposable elements on gene expression responses to abiotic stress in maize, a plant with many families of transposable elements located in between genes. Our analysis suggests that a small number of maize transposable element families may contribute to the response of nearby genes to abiotic stress by providing stress-responsive enhancer-like functions. The specific insertions of transposable elements are often polymorphic within a species. Our data demonstrate that allelic variation for insertions of the transposable elements associated with stress-responsive expression can contribute to variation in the regulation of nearby genes. Thus novel insertions of transposable elements provide a potential mechanism for genes to acquire *cis*-regulatory influences that could contribute to heritable variation for stress response.

## Introduction

Transposable elements (TEs), first described as “controlling elements” by Barbara McClintock [1], are now known to make up the majority of angiosperm DNA [2]-[4]. TE insertions within genes may result in mutant alleles by changing the reading frame or splice pattern, frequently negatively affecting gene function. However, TEs also have the potential to contribute to regulation of gene expression, potentially playing an important role in responses to environmental stress [2], [5]; McClintock initially referred to TEs as “controlling elements” based on their ability to influence the expression of nearby genes [1], [6]. Several specific examples of TE influence on the expression of nearby genes have now been documented (reviewed by [7]-[11]). TE insertions near genes may influence gene expression through several potential mechanisms, including inserting within *cis*-regulatory regions, contributing an outward reading promoter from the TE into the gene [12]-[15], or providing novel *cis*-regulatory sequences that can act as enhancers/ repressors by facilitating transcription factor binding [16], or influencing the chromatin state of gene promoter regions [17]-[19].

Some TEs exhibit stress-responsive transcription or movement [20]-[25]. For example, expression of the tobacco *Tnt1* element can be induced by biotic and abiotic stress [22]-[23]. The rice DNA transposon *mPing* can be activated in response to cold and salt stress [26]-[27]. The Arabidopsis retrotransposon ONSEN is transcriptionally activated by heat stress [16], [28]-[29]. Tissue culture is a complex stress that can result in the activation of DNA transposons in maize and retrotransposons in rice [30]-[31]. There is also evidence that some of these TE responses to environmental conditions can affect the expression of nearby genes. Novel *mPing* MITE insertions in the rice genome in some cases resulted in up-regulation of nearby genes in response to cold or salt stress with no change in expression in control conditions [26]-[27]. The *ONSEN* retrotransposon insertions near Arabidopsis genes exhibit similar properties: alleles containing *ONSEN* insertions often show heat-responsive regulation while alleles lacking *ONSEN* are not up-regulated by heat stress [16]. These studies suggest that TEs can provide novel regulatory mechanisms and influence the response to environmental stress.

Maize provides a good system for studying the potential influence of TEs on regulation of nearby genes. While TEs only account for ∼10% of the Arabidopsis genome [32] or ∼32% of the rice genome [33], they contribute ∼85% to the maize genome [34]-[35]. Many TEs are located in pericentromeric regions and heterochromatic maize knobs [34], [36], but there are also many TE insertions interspersed between maize genes [37]-[39].

The majority of maize genes (66%) are located within 1kb of an annotated transposon [35]. In addition, allelic variation for the presence of TE insertions near genes is high in maize [39] – [41], creating the potential for allelic regulatory differences at nearby genes. For example, polymorphic TE insertions in different haplotypes of the *tb1, Vgt1* and *ZmCCT* loci likely contribute to regulatory differences for these genes [42]-[44].

While there are good examples to suggest that specific TEs can influence the response of nearby genes to abiotic stress [16], [26] it remains unclear how widespread this phenomenon is, how many genes are activated in such a TE-dependent manner, and whether multiple TE families are capable of controlling stress response. We identified a subset of TE families over-represented in the promoters of maize genes that exhibit stress-responsive up-regulation or activation of gene expression. Based on our data, as many as 20% of genes that showed increased expression in response to stress are located near a TE from one of these families. We find that stress-responsive TEs appear to provide enhancer-like activity for nearby promoters and allelic variation for TE insertions is strongly associated with variation in expression response to stress for individual genes.

## Results

We extracted and sequenced RNA from 14 day old seedlings of inbred lines B73, Mo17 and Oh43 grown using standard conditions as well as seedlings that had been subjected to cold (5^0^C for 16 hours), heat (50^0^C for 4 hours), high salt (watered with 300 mM NaCl 20 hours prior to collection) or UV stress (2 hours) (see Materials and Methods for details). For each stress the plants were sampled immediately following the stress treatment and there were no apparent morphological changes in these plants relative to control plants. However, when the stressed plants were allowed to recover for 24 hours under standard conditions phenotypic consequences became apparent for several of the stress treatments (Figure 1A-B). RNAseq data was generated for three biological replicates for cold and heat stress and one sample for the high salt and UV stress (SRA accessions and read number for each sample are provided in Table S1). Differentially expressed genes (RPKM>1 in control or stressed samples, p_adj_<0.1 in DESeq [45] analysis, and minimum of 2-fold change in stress compared to control) were identified in control relative to cold or heat treated plants for each genotype using both the filtered gene set (FGS) and working gene set (WGS) genes (Table S2). For each stress by genotype combination we found that 18%-30% of the expressed genes (7 – 10% of all genes) exhibit significant changes in expression level with similar frequencies of up-and down-regulated expression changes (Table S2). For the salt and UV stress we identified genes that exhibit at least 2-fold change in expression and RPKM >1 in at least one of the conditions. The analysis of data for heat/cold stress revealed that the genes identified as differentially expressed based on a single replicate of this data had >90% overlap with the genes identified as significant in the analysis of multiple replicates. The clustering of gene expression responses to abiotic stress suggests that each stress has a substantial influence on the transcriptome (Figure 1C). While all three inbred lines showed similar transcriptional responses to the stress conditions there is also evidence for genotype-specific responses (Figure 1C).

**Figure 1.**
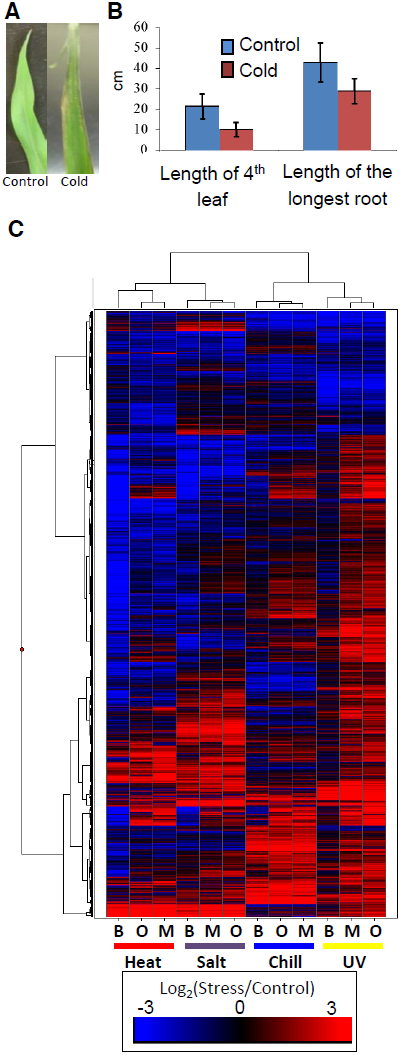
Cold stress effects plant growth and gene expression. (A) Exposure of maize seedlings to cold stress resulted in leaf lesions visible after two days of recovery. A B73 leaf not exposed to cold stress is shown on the left and cold-stressed B73 leaf is shown on the right. **(B)** Seedlings subjected to cold stress showed decreased growth as measured on the 7^th^ day of recovery (p-value < 0.05; 20 plants were measured for each condition; standard error is shown with vertical lines). Similar decreases in growth and fitness were detected for three other stress conditions. **(C)** Abiotic stress exposure results in up-or down-regulation for numerous maize genes in each genotype. The log_2_(stress/ control) values for all differentially expressed FGS genes were used to perform hierarchical clustering of the gene expression values. The genotypes (B73 - B, Mo17 - M, and Oh43 - O) and stress treatments are indicated below each column.

## Some TE families are associated with stress-responsive expression of nearby genes

To test the hypothesis that genes responding to abiotic stress may be influenced by nearby TE insertions we focused our initial analyses on expression responses in the inbred B73, for which a reference genome is available [35]. The TEs located within 1 kb of the transcription start site (TSS) of each gene were identified in the B73 reference genome. For each of 576 annotated TE families we determined whether genes located near the transposon were significantly enriched (p<0.001, >2 fold-enrichment and at least 10 expressed genes associated with the TE family) for responsiveness to each of the stress conditions (separate analyses for enrichment in up- or down-regulated genes for each stress) relative to non-differentially expressed genes (Table S3). While the majority of transposon families are not associated with stress-responsive expression changes for nearby genes (Figure 2A-B; Table S3), 20 TE families are significantly enriched for being located near genes with stress-responsive up-regulation and 3 TE families are associated with genes down-regulated in response to stress (Figure 2C; Table 1).

**Figure 2.**
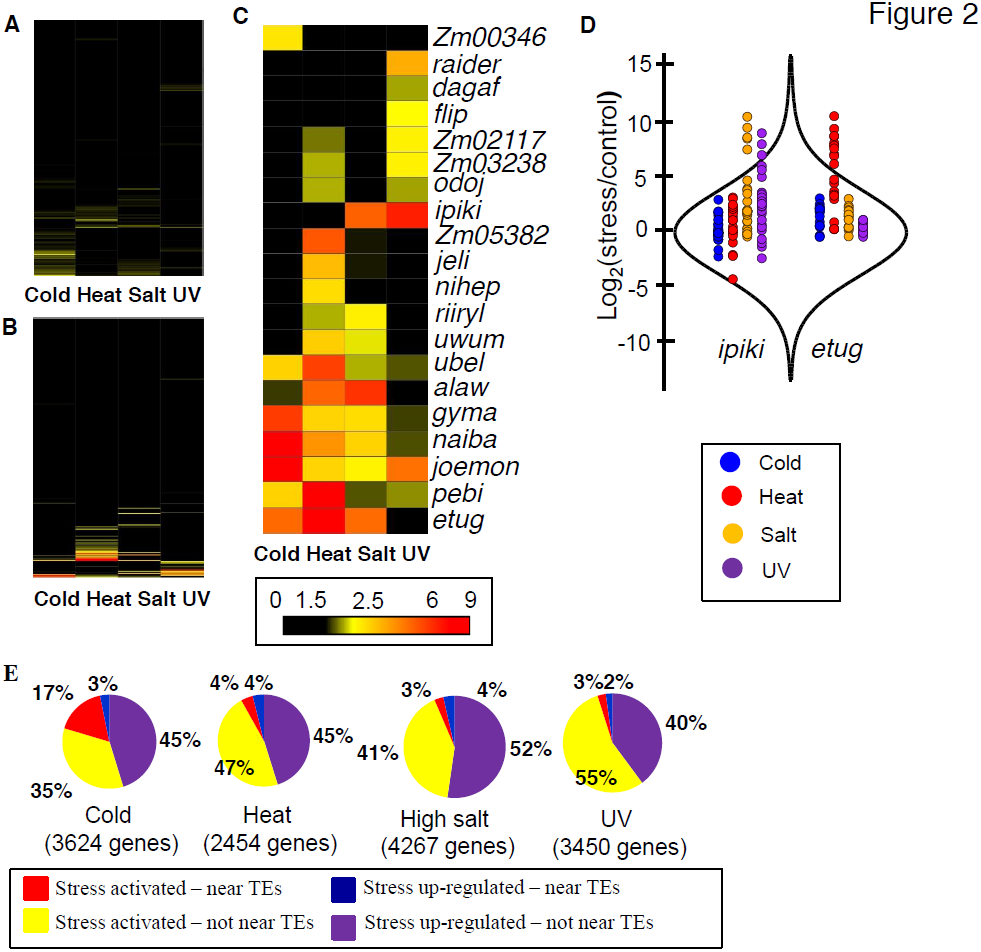
Several TE families are associated with stress-induced up-regulation of gene expression. (A) and (B) Fold enrichment for down-regulated **(A)** and up-regulated **(B)** genes for 283 TE families with the number of expressed WGS genes over 10 is shown as a heat map for four abiotic stress conditions. **(C)** Fold-enrichment values for each of the 20 TE families associated with gene up-regulation in response to abiotic stress are shown as a heat map. (**D**) Comparison of distributions of log_2_ (stress/control) values between all genes and genes located near certain TE families. The distribution of all genes is shown using a violin plot while the expression changes for individual genes are shown using colored dots. Genes located near *ipiki* elements are shown on the left and genes located near *etug* elements are shown on the right with the colors indicating the different environmental stresses. **(E)** The relative proportion of WGS genes turned on or up-regulated following stress that are associated with the TE families (from **C**) is indicated for each stress condition in B73. Total number of up-regulated genes is shown for each stress. The expected proportion of genes with insertions of TEs from the enriched families for all expressed genes is less than 1% for all stresses.

**Table 1.**
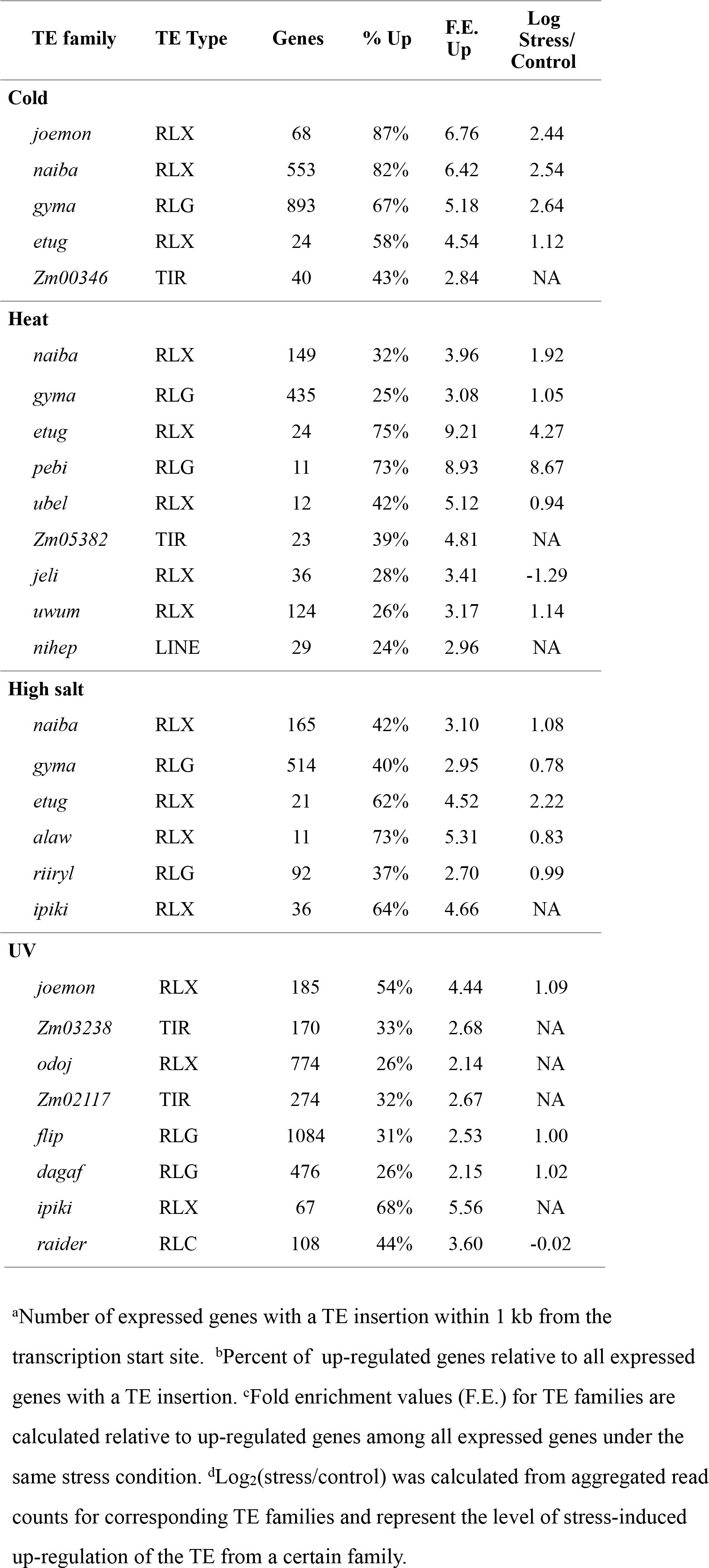
TE families enriched for genes up-regulated in response to abiotic stress.

Examples of the expression changes for genes in different abiotic stresses are shown for two transposon families, *ipiki* and *etug* (Figure 2D). Genes located near *ipiki* are enriched for up-regulation following salt and UV stress while genes located near *etug* elements are enriched for heat-responsive up-regulation. One striking example is the *joemon* TE family for which 59 of 68 expressed genes containing an insertion within 1 kb are activated following cold stress (Table 1). Although similar numbers of genes exhibit increased and decreased gene expression genome-wide following abiotic stress conditions, the majority of enriched TE family – stress combinations (28/31) are associated with up-regulated gene expression. For each of the stress conditions there were 4-9 TE families that are associated with up-regulation of gene expression. Some TE families are associated with altered expression in multiple stress treatments (Table 1, Table S4; Figure 2C) and two of the TE families associated with down-regulation of gene expression under high salt stress were also associated with increased gene expression under UV stress.

The TE families enriched for genes activated in response to stress include all major super-families of TEs: TIR DNA transposons, LTR *gypsy*-like (RLG), *copia*-like (RLC), or unknown (RLX) retrotransposons, and LINE elements (Table 1,). These TE families vary substantially for the number of genes that they are located near: from 30 to 3052 genes (Table 1; Table S4) and are spread uniformly across the maize genome. The presence of these TEs near genes is not fully sufficient for stress-responsive expression.

For each of the TE families identified, 26 – 87% of genes located near a TE insertion show stress responsive expression depending on the stress and the TE family. The expression levels for the TEs themselves was assessed for each of the treatments and in the majority of TE family – stress combinations (14 of 21 with expression data) the TEs showed at least 2-fold increase in transcript levels in the stress treatment compared to control conditions (Table 1, Table S4). There are several examples of TE families that exhibit increased levels of expression in a particular stress but the nearby genes are not enriched for stress-responsive expression (Table S3), suggesting that not all TEs that are influenced by a particular stress influence nearby genes.

To understand what proportion of the transcriptome response to a specific abiotic stress may be explained by influences of specific TEs inserted near genes, up-regulated genes were classified according to whether they were located near a member of one of the stress-associated TE families (1 kb 5’ from TSS) and whether they are up-regulated (expressed under control and stress conditions) or activated in response to stress (only expressed following stress treatment). We found that a substantial portion of the transcriptome response to the abiotic stress could be associated with genes located near the set of 4-9 TE families that were identified as enriched for up-regulated genes (Figure 2E). In total, 5-20% of the genome-wide transcriptome response to the abiotic stress and as many as 33% of activated genes could be attributed to the genes located near one of these TE families (Figure 2E; Table S5-6).

## Some TE families act as local enhancers of stress-responsive expression

One possible mechanism by which these families of TEs could contribute to stress-responsive expression for nearby genes is that the TE may provide an outward-reading promoter that is stress-responsive. This model predicts that the orientation of the TE relative to the gene is important and that novel transcripts containing TE sequences fused to gene sequences would be present for up-regulated genes under stress conditions. In order to assess the importance of the orientation of the TE insertion relative to the gene, we compared the proportion of genes located on the same strand as a TE for genes up-regulated in response to stress and genes non-differentially expressed in response to stress for all TE families enriched for up-regulated genes (Table S7). While most families showed no significant difference in the proportion of genes on the same strand as the TE between the up-regulated and non-differentially expressed genes, a minority of families (4/20) showed significant enrichment. For example, 97% of the stress-responsive genes located near *etug* elements are on the same strand as the TE (Table S7). Nonetheless, visual inspection of the RNAseq alignments did not reveal evidence for stress-responsive transcripts that initiate within the TE and include the gene.

Alternative models include the possibility that the TE may contain *cis*-regulatory sequences that can act as binding sites for stress-induced transcription factors, or that the TE could influence the local chromatin environment in such a way that the region is more accessible under stress conditions. The analysis of TE distance from transcription start sites of stress-responsive genes suggests that in many cases the effect of TE on stress-responsive gene activation quickly diminishes as the distance increases beyond 500 bp – 1kb (Figure S1A). The DREB/CBF transcription factors are often involved in transcriptional responses to abiotic stress in plants [46]. The consensus sequence for DREB/CBF binding (A/GCCGACNT [47]) was found in most of the TEs that were associated with stress-responsive expression for nearby genes, with the exception of elements that only exhibit UV stress response (Figure S1B). While we did not have evidence to distinguish between the possibilities that TEs provide either a sequence-specific binding site that might act as a stress-specific enhancer or influence the chromatin state in a non-sequence specific manner, our data are consistent with the TE insertions acting predominantly as local enhancers of expression rather than as novel promoters.

Because individual TE copies are subject to frequent rearrangements and internal deletions, we investigated whether the presence of specific regions in each TE family were over-represented in insertions that confer stress-responsive expression. For six of the 20 TE families, this comparison revealed specific portions of the TE sequences enriched among insertions that convey stress-responsive expression. For example, *naiba* and *etug* insertions located near up-regulated genes are approximately four times as likely to contain a particular portion of the TE long terminal repeat (LTR; p-value < 0.001; Fig. S2), and this same sequence is found in a subset of insertions of the related family, *gyma,* that are associated with up-regulated genes. While we did not have evidence to rule out the possibility that TEs influence the chromatin state in a non-sequence specific manner, these data indicate that the presence of particular regions of TE elements likely provide enhancer functions associated with gene expression responses to stress and help explain the variable effect of different insertions of the same family on stress-responsive expression.

## Characterization of genes with TE-influenced stress responsive expression

We assessed a number of properties of the TE-influenced stress-responsive genes in comparison with stress-responsive genes that are not associated with one of these TE families (Table 2). Stress-responsive genes located near the TE families tend to be substantially shorter in length with fewer introns. Analysis of developmental expression patterns for these genes using the B73 expression atlas [48] reveals that only 7% of the TE influenced genes are expressed in at least 5 tissues, compared to 41% of the non-TE influenced genes. The TE influenced genes are also less likely to be in the filtered gene set (FGS), and the proportion of the TE influenced genes with syntenic homologs in other grass species is much lower than the proportion of non-TE influenced genes (Table 2). Each of these features was assessed separately for each of the TE families (Table S7) and there is some variation for these properties among different families. These observations are compatible with the notion that TE insertions may in some cases function as enhancers that can drive expression of cryptic promoters in non-coding regions of the genome. This will result in stress-responsive production of transcripts that may be annotated as genes but may not produce functional proteins. However, 37% of TE influenced genes are included in the FGS that has been curated to remove transposon-derived sequences and a substantial proportion of the TE influenced genes are syntenic with genes from other species, have GO annotations, and could contribute to functional responses to stress (Table 2, S7). These results suggests that many of TE influenced genes are not derived from TEs.

**Table 2.**
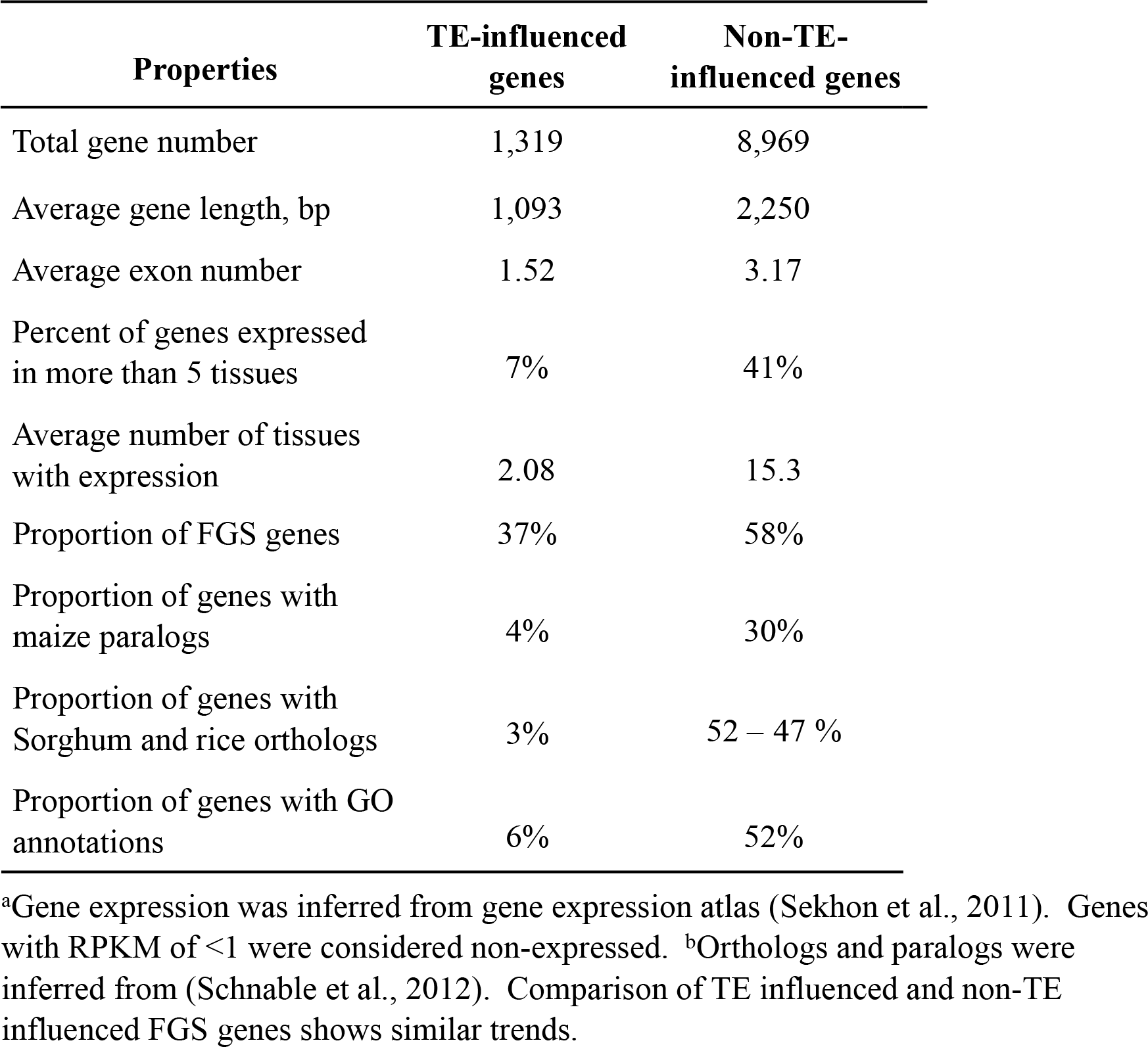
Comparison of TE-influenced and non-TE-influenced WGS genes up-regulated in abiotic stress

## Contribution of TEs to allelic variation for stress-responsive expression

We were particularly intrigued by the question of whether polymorphic insertions of TEs from families associated with stress-responsive expression of nearby genes might contribute to allelic variation for stress-responsive gene expression. The consistency of stress-responsive expression of TE-associated genes across the three inbred lines surveyed varied widely across TE families (Figure 3A; Figure S3). In order to assess whether insertions of TEs from the families associated with stress-responsive gene expression could contribute to allelic variation for gene expression regulation, we used whole-genome shotgun re-sequencing data from Mo17 and Oh43 [49] to find potential novel insertions of elements from the TE families identified in this study. We identified 23 novel (not present in B73) high-confidence insertions of TEs from these families located within 1kb of the TSS of maize genes and validated them by PCR (Table S8). Of the 10 genes with detectable expression in our RNAseq experiments, 7 showed stress-responsive up-regulation / activation associated with the TE-containing alleles (Figure 3B). This analysis was expanded to additional genotypes by using PCR to detect the presence/absence of the TE insertion in a diverse set of 29 maize inbred lines. The relative expression of the gene in stress compared to control treatment was also determined in each inbred using quantitative RT-PCR. For each of these genes we found that the alleles that lack the transposon insertion did not exhibit stress-responsive expression (Figure 4), with the exception of one genotype for gene GRMZM2G108057. In contrast, the majority of the alleles that contain the TE (60-88%) exhibit stress-responsive up-regulation. Although for a single insertion we cannot rule out the possibility that differential expression is due to a different polymorphism on the same haplotype as the TE, the fact that we see TE-associated expression change in multiple genes for each of the TE families (Table. S8) argues strongly against such an explanation in general. These data thus provide evidence that insertion polymorphisms for the TE families identified here can generate novel expression responses for nearby genes.

**Figure 3.**
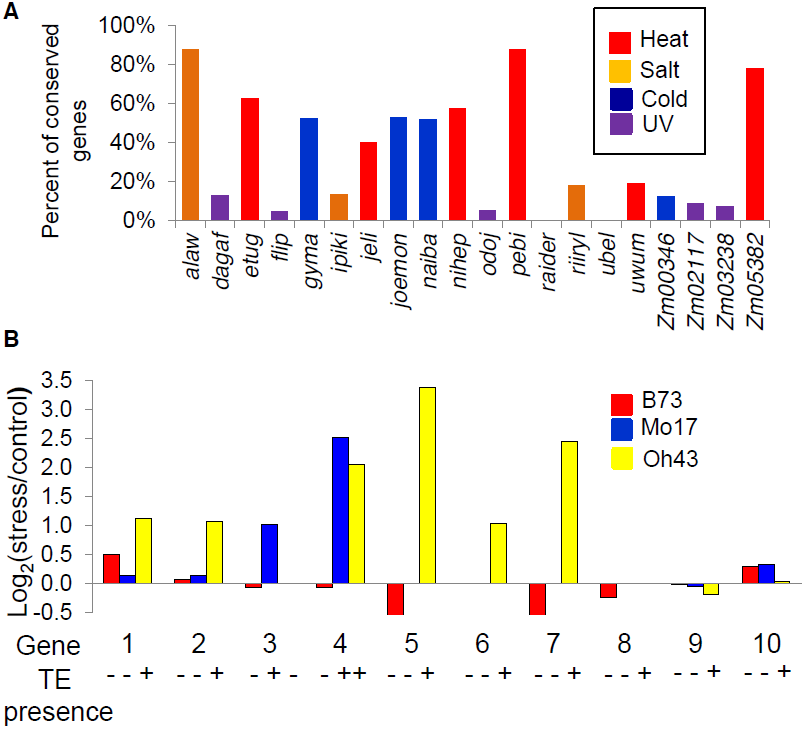
Stress-induced up-regulation of gene expression correlates with the variation in TE presence. (A) Proportion of genes up-regulated in B73 that are also up-regulated in Mo17 and Oh43 is shown for all TE families under the stress condition with highest enrichment for the TE family. **(B)** The relative expression levels in stress compared to control treatments (log_2_ ratio) is shown for B73, Mo17, and Oh43 for each of the 10 expressed genes that are polymorphic for insertions of TEs. The presence/ absence of the TE for each genotype-inbred combination is shown by ‘+’ and ‘-‘ symbols. The genes are as follows: 1-GRMZM2G102447; 2-GRMZM2G108057; 3-GRMZM2G071206; 4-GRMZM2G108149; 5-GRMZM2G400718; 6-GRMZM2G347899; 7-GRMZM2G517127; 8-GRMZM2G378770; 9-GRMZM2G177923; 10-GRMZM2G504524. All genes with TE insertion polymorphism are listed in Table S8.

**Figure 4.**
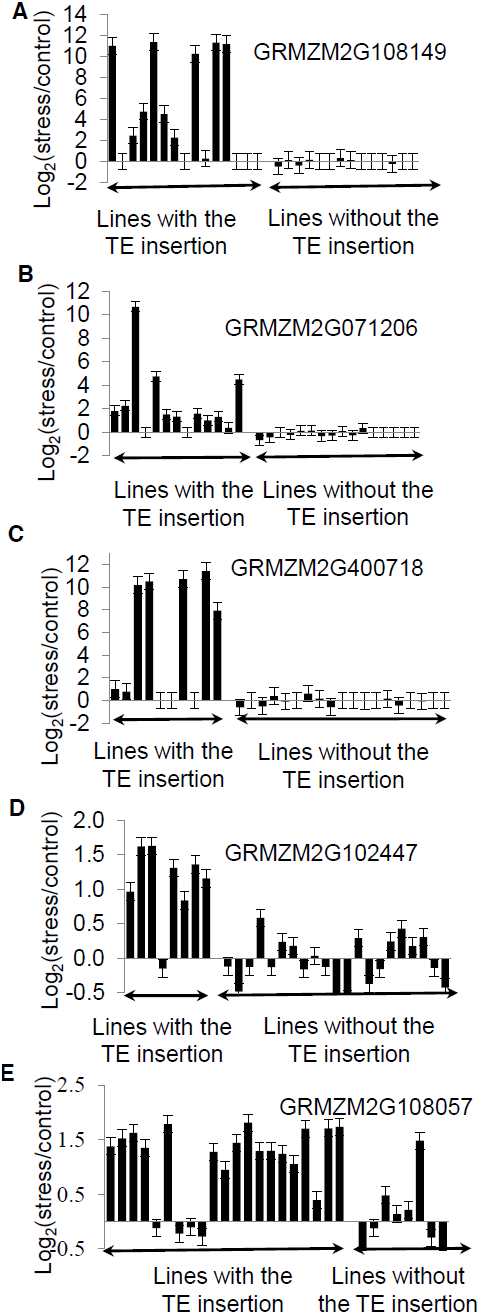
Validations of correlation between stress-induced up-regulation of gene expression and presence of TEs. The presence / absence of insertions of ZM00346 elements in the promoter of GRMZM2G108149 **(A)**, GRMZM2G071206 **(B)**, GRMZM2G400718 **(C)**, GRMZM2G102447 **(D)**, and GRMZM2G108057 **(E)** was assessed by PCR and genotypes were divided according to whether this insertion is present or not (displayed in alphabetical order). The changes in gene expression are shown as log_2_(stress/control) values determined using qRT-PCR for each genotype. Vertical brackets correspond to standard error based on three technical replicates of qRT-PCR experiments.

## Discussion

Transposable elements are a major component of many eukaryotic genomes, and constitute the majority of plant nuclear DNA. TEs are usually considered as a deleterious or neutral component of these genomes. However, the interplay between TEs and genes may have important functional contributions to plant traits. There are clear examples of TE insertions that are linked to functionally relevant alleles in maize such as *Tb1* [42] *Vgt1* [43] *and ZmCCT* [44]. In these cases, a transposon insertion within a distant cis-regulatory sequence influences the regulation of adjacent genes. There are also examples of functionally relevant TE insertions in tomato, melons and citrus [50]-[52] that can influence gene expression, potentially through chromatin influences that generate obligate epialleles.

Previous research in several plant species has suggested that at least some families of transposable elements may become transcriptionally activated following environmental stress. Tissue culture has been shown to result in activation of transposons and retrotransposons in a number of plant species [30]-[31]. There are also examples of transcriptional activation of TEs in response to specific abiotic stresses in tobacco [22], rice [26]-[27] and Arabidopsis [16], [28]-[29]. It is expected that the stress responsive expression of these TEs involves local enhancers that result in up-regulation of the TE promoter in response to stress. These local enhancers could also act upon other nearby promoters. There are a handful of examples in which transposon insertions have been linked to stress-responsive expression of nearby genes including the mPING insertions associated with cold-responsive expression in rice [26]-[27] and ONSEN insertions associated with heat-stress responsive expression in Arabidopsis [16]. If this is a common occurrence then we might expect it to be even more prevalent in a genome such as maize where many genes are closely surrounded by TEs.

Our analysis suggested that a small number of TE families are associated with stress-responsive expression for nearby genes. While some TE families were associated with multiple stresses, we found a different subset of TE families for each abiotic stress that was evaluated. In most cases, these same TEs themselves were up-regulated in response to the stress treatment. However, we also noted that there were some TE families that themselves exhibit strong up-regulation but did not have apparent influences on a significant portion of nearby genes. Even though the majority of stress responsive regulation of gene expression is not associated with TEs, based on our data, up to 20% of genes up-regulated in response to stress and as many as 33% of genes activated in response to stress could be attributed to regulation by TEs. One of the alternative explanations would argue that only a small number of genes localized close to a TE are truly influenced by this TE insertion for their expression, while other up-regulated genes are secondary targets and are regulated by the TE influenced genes. Although some of the TE influenced genes we identified could be secondary targets, secondary target genes would not preferentially co-localize with TEs from specific families.

The analysis of the nearby genes that were influenced by TEs suggests that many of them may not actually be protein coding genes. In one sense, this is an expected result. If an enhancer sequence is mobilized within the genome it will have the potential to influence expression from both gene promoter as well as cryptic promoters that may not be associated with coding sequences. The gene annotation efforts in maize have relied upon EST and RNA-seq expression data from a variety of conditions. In many cases the genes that were found to exhibit stress-responsive expression associated with TEs were only annotated as genes based upon evidence of their expression. We would expect that insertions of the TEs that provide stress-responsive enhancer activity would influence cryptic promoters not associated with genes in many cases, but would also affect the expression of nearby protein coding genes. The frequency of each appeared to vary among TE families, with some, like *nihep,* showing little difference between TE-influenced and non-TE-influenced up-regulated genes (Table S7). Overall, while TE influenced stress-responsive genes are enriched for short sequences with limited homology to sequences in other species, a significant proportion are longer, have several exons, are conserved in other species, and have GO annotations.

A particularly interesting aspect of these results is the potential mechanism for creating novel *cis*-regulatory variation. Our understanding of how particular genes might acquire novel regulatory mechanisms is limited. In many cases SNPs within promoters or regulatory sequences have limited functional significance. Therefore, it is difficult to envision how a novel response to a particular environmental or developmental cue would arise. Variation in TE insertions has the potential to create novel regulatory alleles by providing binding sites for transcription factors or influencing chromatin. We provide evidence that allelic variation for stress-responsive expression can be created by the insertion of certain TEs. Variation in TE insertions would generate allelic diversity that could influence an organism's response to environmental conditions and would provide phenotypic variation that could be acted upon by selection. As with other types of variation, most examples of novel stress-responsive expression are likely to be neutral or deleterious and would not be expected to rise in allele frequency. However, a subset of novel stress-responsive expression patterns could be beneficial and become targets of natural or artificial selection contributing to gene regulation networks of environmental stress response.

## Materials and Methods

### Plant growth and stress conditions

B73, Mo17, and Oh43 maize seedlings were grown at 24^0^C in 1:1 mix of autoclaved field soil and MetroMix under natural light conditions in July 2013. For cold stress, seedlings were incubated at 5^0^C for 16 hours. For heat stress, seedlings were incubated at 50^0^C for 4 hours. For high salt stress, plants were watered with 300 mM NaCl 20 hours prior to tissue collection. UV stress was applied in the growth chamber conditions using UV-B lamps for 2 hours prior to tissue collection. UV stress causes accumulation of DNA mutations but most of such mutations would either have no immediate effect on gene expression or would lead to decrease or abortion of expression of specific genes. Light conditions were the same for all stress and control conditions. Whole above ground tissue was collected for 14 day old seedlings at 9am and six seedlings were pooled together for each sample. Three replicates for heat and cold-treated B73 and Mo17 seedlings were grown 3 days apart.

### RNA isolation and RNAseq analysis

Three biological replicates of cold and heat stress and control conditions for B73 and Mo17 were prepared with eight plants pooled for each of the replicates. One biological replicate of high salt and UV stress conditions for B73 and Mo17 as well as all four stress and control conditions for Oh43 were prepared similarly. RNA was isolated using Trizol (Life Technologies, NY, USA) and purified with LiCl. All RNA samples were prepared by the University of Minnesota BioMedical Genomics Center in accordance with the TruSeq library creation protocol (Illumina, San Diego, CA). Samples were sequenced on the HiSeq 2000 developing 10-20 million reads per sample. Transcript abundance was calculated by mapping reads to the combined transcript models of the maize reference genome (AGPv2) using TopHat [53]. Reads were filtered to allow for only uniquely mapped reads. A high degree of correlation between replicates was observed (r>0.98). RPKM values were developed using ‘BAM to Counts’ across the exon space of the maize genome reference working gene set (ZmB73_5a) within the iPlant Discovery Environment (www.iplantcollaborative.org). Genes were considered to be expressed if RPKM>1 and differentially expressed if log_2_(stress/control) > 1 or log_2_(stress/control) < −1. Statistical significance of expression differences was determined using DeSeq package for all fully replicated samples [45].

### Data Analysis

For each gene, transposons located within 1 kb of the transcription start site (TSS) were identified using the B73 reference genome annotation [35] and maize TE elements database [34]. TE distance from transcription start sites was determined using the *closestBed* tool from the BEDTools suite [54] where TEs upstream were given a positive distance value and TEs downstream were given a negative distance value. The transcriptional start site was defined as the 100-bp window intersecting the first base pair of a gene model from the maize genome gene set (ZmB73_5b). The proportion of up-regulated, down-regulated, and non-differentially expressed genes that have an insertion of a TE element from a particular family was calculated for 576 TE families for four stress conditions. Fold-enrichment of up-regulated genes relative to all expressed genes (the sum of up-regulated, down-regulated and non-differentially expressed genes) and relative to all genes was calculated for all TE family / stress combinations. Given the total number of expressed genes associated with each TE family and the proportion of up- and down-regulated genes, the expected numbers of up- and down-regulated genes and non-differentially expressed genes were calculated and a multinomial fit test was conducted. TE families that had over 10 expressed genes associated with them, fold enrichment of up- or down-regulated genes over 2, and p value <0.001 were considered “enriched” for up- or down-regulated, respectively. Similar analysis was conducted for working gene set and filtered gene set genes. The same set of “enriched” TE families was found for both groups of genes as well as when fold enrichment was calculated relative to all expressed genes or to all genes associated with TEs from a particular family.

To assess expression changes in response to stress for TE families, the *overlap* tool from BEDTools suite [54] was used to obtain read counts per each TE accession. The output file from alignment (BAM) was mapped to TE positions listed in the TE GFF file downloaded from maizesequence.org. Each read was required to have 100% overlap with a given TE region. The reads mapping to more than 5 locations in the genome were omitted. The reads were then summed across the entire TE region and combined for each of the TE families.

Tissue specific expression data is from the maize gene expression atlas [47]. Genes with RPKM of <1 were considered non-expressed. Orthologous and paralogous gene pairs were inferred from [55].

### TE polymorphism prediction and verification

Nonreference TE insertions were detected for Oh43 and Mo17 using relocaTE [56], whole genome sequence from the NCBI SRA (Oh43: SRR447831-SRR447847; Mo17: SRR447948-SRR447950), and consensus TE sequences from the maize TE database [34]. Reads containing TEs were identified by mapping to consensus TE sequences, trimming portions of reads mapping to a TE, and mapping the remaining sequence to the reference genome. Nonreference TEs were identified when at least one uniquely mapped read supported both flanking sequences of the nonreference TE, overlapping for a characteristic distance that reflects the target site duplication generated upon integration (five nucleotides for all LTR retrotransposons, nine nucleotides for DNA TIR mutator). Primers for six TE polymorphic genes up-regulated under stress conditions in Oh43 or Mo17 but not in B73 were designed using Primer 3.0 software [57] and PCR reactions were performed using Hot Start Taq Polymerase (Qiagen, Ca, USA). Primer sequences are shown in Supplementary Table 10.

### cDNA synthesis and qPCR

cDNA synthesis and qPCR analysis were performed as described in [58]. Primers for 10 differentially expressed genes and two control genes (*GAPC* and *mez1*) were designed using Primer 3.0 software [57]. Primer sequences are shown in Supplementary Table 10.

## Acknowledgements

The authors are grateful to Peter Hermanson, Jaclyn Noshay, Hailey Karlovich, Josie Slater, Amanda Nimis, and Kristin Male for help in developing protocols for stress conditions, collecting samples, and data validation.

## Supporting Information

**Figure S1.**
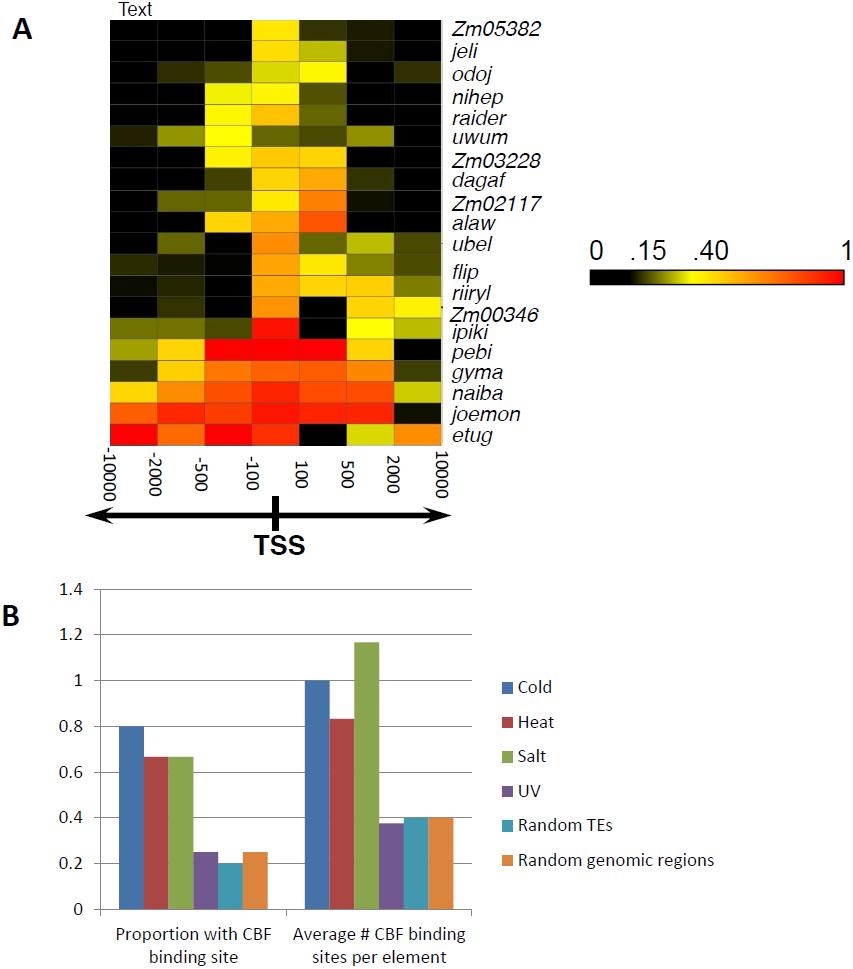
Properties of TE insertions that condition stress-responsive expression. (A) In our initial screening we only analyzed TE insertions located within 1kb of the TSS. Here we assessed the proportion of genes that exhibit stress-responsive expression for TE insertions located at different distances from the TSS (for the stress condition most associated with each TE family). Some of the TE families appear to only affect genes if they are inserted quite near the TSS while others can have influences at distances. (B) The CBF/DREB transcription factors have been associated with stress-responsive expression in a number of plant species [46]. We identified consensus CBF/DREB binding sites (A/GCCGACNT) in the consensus TE sequences (maizetedb.org) for the TEs associated with each of the stresses as well as in 40 randomly selected TEs that were not associated with gene expression responses to stress or 40 randomly selected 5kb genomic regions. The proportion of sequences that contained a CBF/DREB binding site and the average number of sites per element are shown. The TEs associated with cold, heat and salt stress are all enriched for containing CBF/DREB binding sites.

**Figure S2.**
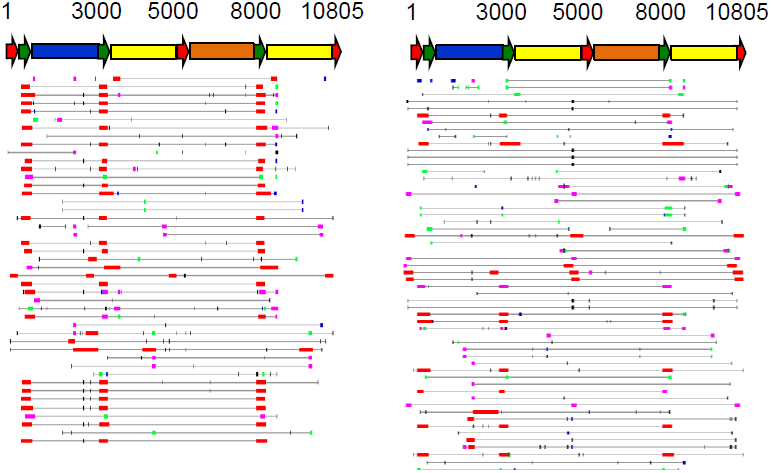
TE insertions co-localized with TE-influenced stress-responsive genes frequently share the same part of the TE element. Alignment of unique *naiba* insertions co-localized with cold-responsive (left) and stress-non-responsive (right) genes are shown. *Naiba* element structure is shown on top with various colors representing repeated regions of the element. The region that differentiates mostly between up-regulated and non-differentially expressed genes is a repeat and is shown as a green arrow. The same sequence is shared by a subset of *gyma* elements co-localized with up-regulated genes.

**Figure S3.**
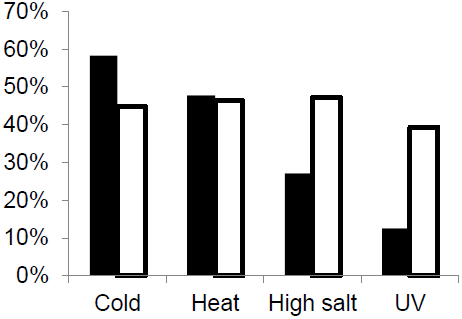
The conservation of stress-responsive expression of TE influenced genes varies for different families and different stresses. Proportion of genes up-regulated in B73 that are also up-regulated in Mo17 and Oh43 is shown for all four stresses for TE-influenced (black) and non-TE influenced (white) genes.

**Table S1.** Sequencing depth for the samples used in this study.

**Table S2.** Gene expression response to abiotic stress in maize seedlings.

**Table S3.** Relationships between genes affected by abiotic stress and TE elements located within 1000 bp of a gene transcription start site.

**Table S4.** TE families enriched for genes up-regulated in response to abiotic stress. **Table S5.** List of TE influenced and non-TE influenced genes activated in response to abiotic stress.

**Table S6.** Number of TE influenced and non-TE influenced genes up-regulated in response to abiotic stress.

**Table S7.** Characteristics of TE families enriched for genes up-regulated by abiotic stress. **Table S8.** Validation of stress-induced activation of genes located near novel TE insertions in Oh43 and Mo17.

**Table S9.** Validation of associations between TE polymorphisms and stress-induced gene activation in diverse inbred lines.

**Table S10.** List of primers used in the study.

